# Mechanisms underpinning morphogenesis of symbiotic organ specialized for hosting indispensable microbial symbiont in stinkbug

**DOI:** 10.1101/2023.03.02.530924

**Authors:** Sayumi Oishi, Toshiyuki Harumoto, Keiko Okamoto-Furuta, Minoru Moriyama, Takema Fukatsu

## Abstract

Microbial mutualists are pivotal for insect adaptation, which often entails the evolution of elaborate organs for symbiosis. Addressing what mechanisms underpin the development of such organs is of evolutionary interest. Here we investigated the stinkbug *Plautia stali* whose posterior midgut is transformed into a specialized symbiotic organ. Despite being a simple tube in newborns, it developed numerous crypts in four rows, whose inner cavity hosts a specific bacterial symbiont, during 1^st^ to 2^nd^ nymphal instar. Visualization of dividing cells revealed that active cell proliferation was coincident with the crypt formation, although spatial patterns of the proliferating cells did not reflect the crypt arrangement. Visualization of visceral muscles in the midgut, consisting of circular muscles and longitudinal muscles, uncovered that, strikingly, circular muscles exhibited a characteristic arrangement running between the crypts specifically in the symbiotic organ. Even in early 1^st^ instar when no crypts were seen, two rows of epithelial areas delineated by bifurcated circular muscles were identified. In 2^nd^ instar, crossing muscle fibers newly appeared and connected the adjacent circular muscles, whereby the midgut epithelium was divided into four rows of crypt-to-be areas. The crypt formation proceeded even in aposymbiotic nymphs, uncovering autonomous nature of the crypt development. We propose a mechanistic model of crypt formation wherein the spatial arrangement of muscle fibers and the proliferation of epithelial cells underpin the formation of crypts as midgut evaginations.

**IMPORTANCE:** Diverse organisms are associated with microbial mutualists, in which specialized host organs often develop for retaining the microbial partners. In the light of the origin of evolutionary novelties, it is important to understand what mechanisms underpin the elaborate morphogenesis of such symbiotic organs, which must have been shaped through interactions with the microbial symbionts. Using the stinkbug *Plautia stali* as a model, we demonstrated that visceral muscular patterning and proliferation of intestinal epithelial cells during early nymphal stages are involved in the formation of numerous symbiont-harboring crypts arranged in four rows in the posterior midgut to constitute the symbiotic organ. Strikingly, the crypt formation occurred normally even in symbiont-free nymphs, uncovering that the crypt development proceeds autonomously. These findings suggest that the crypt formation is deeply implemented into the normal development of *P. stali*, which must reflect the considerably ancient evolutionary origin of the midgut symbiotic organ in stinkbugs.

## INTRODUCTION

Diverse insects are obligatorily associated with microbial mutualists (1, 2). The microbial partners usually play important biological roles for their insect hosts, which encompass nutrient provisioning (3), food digestion (4), defense against enemies (5), tolerance to abiotic stresses (6) and others. For hosting the microbial partners, the insects often develop specialized “symbiotic organs” (1, 7). In aphids, for example, their essential bacterial symbionts are endocellularly harbored in specialized cells and organs for symbiosis, so-called the bacteriocytes and the bacteriomes (8). In stinkbugs, their essential bacterial symbionts are extracellularly harbored in gut-associated symbiotic organs with sac- or pouch-like structures, so-called the crypts or the gastric caeca (9).

The evolutionary origin of the symbiotic organs has been regarded as a challenging evo-devo issue in the light of the origin of evolutionary novelties (10–12). As for molecular mechanisms underpinning the embryonic differentiation and development of the symbiotic organs, co-option of Hox transcription factors has been identified in aphids, seed bugs and ants (13–15). In the postembryonic development, the symbiotic organs often exhibit remarkable morphological traits (1), but it has been poorly understood what mechanisms govern the spectacular morphogenesis of the symbiotic organs.

In this context, the brown-winged green stinkbug *Plautia stali* (Hemiptera: Pentatomidae) (**Fig. 1a**) provides an experimentally tractable model symbiotic system (16, 17). The midgut of *P. stali* is differentiated into structurally distinct M1 to M4 regions (**Fig. 1b-d**), of which the most posterior M4 region is specialized as a symbiotic organ with numerous crypts arranged in four rows (**Fig. 1e, f**). The inner cavity of each pouch-like crypt harbors the γ-proteobacterial symbiont *Pantoea* sp., which is essential for normal growth and survival of the host insects and vertically transmitted to newborns via egg surface contamination (16, 18). Our previous study described the developmental process of the symbiotic organ of *P. stali* in detail morphologically and histologically (19), which unveiled that, although the posterior midgut is a simple tube upon hatching, it develops numerous crypts in four rows during 1^st^ to 2^nd^ nymphal instar (**Fig. 1g-j**). The crypt formation in the posterior midgut must be important for the stinkbug-microbe mutualism in that the structural configuration of the numerous crypts enables stable retention of the symbiotic bacteria and also increases the epithelial surface area for host-symbiont metabolic interactions. However, it has been elusive what mechanisms underlie the formation of the highly ordered arrangement of the crypts in the midgut symbiotic organ.

**FIG 1.**
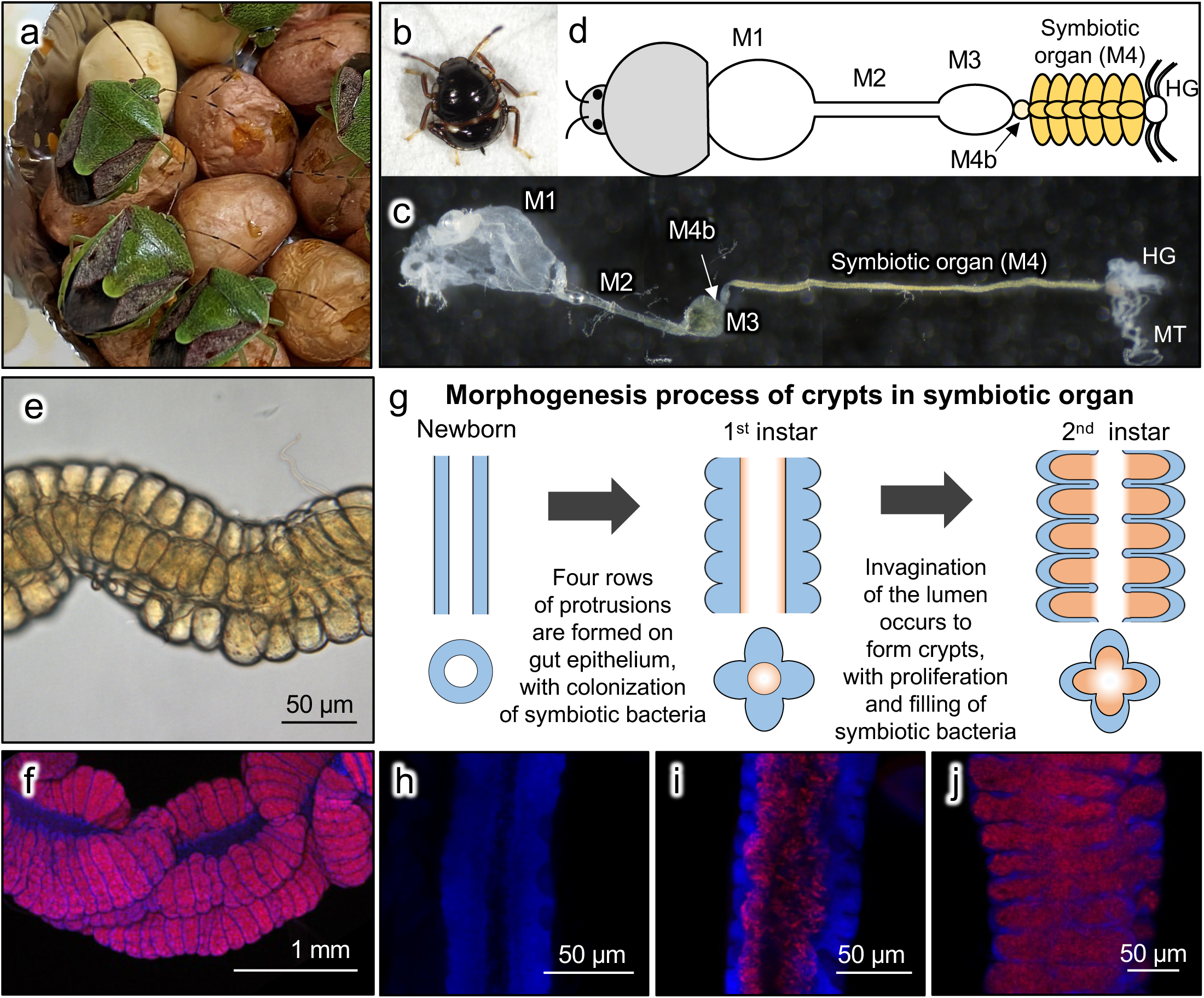
Arrangement and formation process of the crypt rows of the midgut symbiotic organ on the alimentary tract of *P. stali*. (a) Adult insects reared on peanuts and soybeans. (b) A 2^nd^ instar nymph. (c, d) A dissected alimentary tract of the 2^nd^ instar nymph (c) and its schematic representation (d). Abbreviations: M1, midgut 1^st^ section; M2, midgut 2^nd^ section; M3, midgut 3^rd^ section; M4b, bulb-like midgut section anterior to M4; M4, midgut 4^th^ section with crypts (symbiotic organ); HG, hindgut; MT, Malpighian tubule. (e) A microscopic image of the M4 symbiotic region, on which 3 of 4 crypt rows are seen with a crypt row hidden behind. (f) FISH of adult’s symbiotic organ. (g) Schematic diagrams of the crypt morphogenesis in the symbiotic organ. Also see (19). Blue and orange indicate the host intestinal epithelium and the symbiotic bacteria, respectively. (h) FISH of symbiotic organ of newborn nymph. (i) FISH of symbiotic organ of 1^st^ instar nymph 1 day after hatching. (j) FISH of symbiotic organ of 2^nd^ instar nymph 1 day after molting. In (f, h-j), red indicates symbiotic bacteria and blue indicates cell nuclei, respectively.

In this study, we investigated the developmental and morphogenetic processes of the posterior midgut region of *P. stali* in detail, particularly focusing on 1^st^ and 2^nd^ nymphal instars when the crypt formation occurs. We uncovered that stage-specific proliferation of gut epithelial cells and characteristic spatial arrangement of visceral muscle fibers are involved in the crypt morphogenesis, which proceeds autonomously even in the absence of the symbiotic bacteria.

## RESULTS

### Cell proliferation patterns in the nymphal symbiotic organ during crypt formation of *P. stali*

First, we investigated the cell proliferation patterns in the posterior midgut region M4, where crypt morphogenesis occurs to form the symbiotic organ, during the 1^st^ and 2^nd^ nymphal instars of *P. stali* (see **Fig. 1g**). In the developmental course of the 1^st^ and 2^nd^ instar nymphs, we visualized DNA synthesizing cells by EdU labeling and dividing cells by H3P antibody staining, respectively, in the symbiotic organ. Both DNA synthesizing cells and dividing cells were scarce in the early 1^st^ instar, steadily increased toward the mid to late 1^st^ instar, and became scarce again in the 2^nd^ instar (**Fig. 2**). It should be noted that the crypt morphogenesis starts and proceeds in the mid to late 1^st^ instar period (see **Fig. 1g**) (19). These observations suggested that the crypt formation in the mid to late 1^st^ instar nymphs entails activated cell proliferation in the symbiotic organ. On the other hand, neither the DNA synthesizing cells nor the dividing cells exhibited specific patterns corresponding to the crypt structures (**Fig. 2h-j**), which suggested that the cell proliferation alone cannot account for the formation of the characteristic crypt patterns.

**FIG 2.**
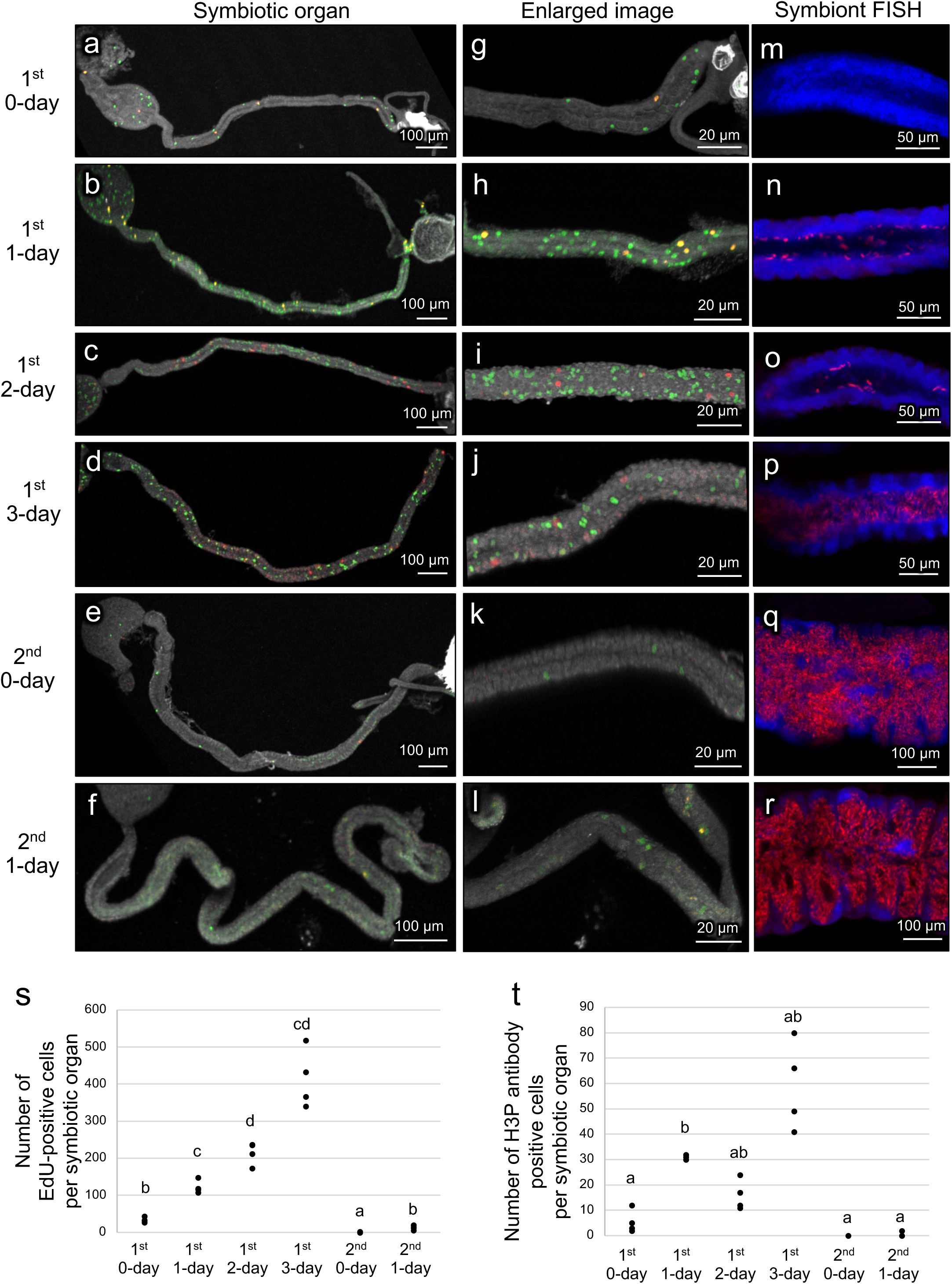
Cell proliferation dynamics of the symbiotic organ in the nymphal development of *P. stali*. (a-f) Visualization of DNA synthesizing cells by EdU labeling (green), dividing cells by H3P antibody staining (red), and all intestinal cells by DNA staining (white) in dissected midgut preparations. (g-l) Enlarged images corresponding to (a-f) at the midgut symbiotic region where crypts are formed. (m-r) FISH of symbiotic organ. Red indicates symbiotic bacteria and blue indicates cell nuclei, respectively. (a, g, m) 1^st^ instar nymph 0-day after hatching. (b, h, n) 1^st^ instar nymph 1-day after hatching. (c, i, o) 1^st^ instar nymph 2-day after hatching. (d, j, p) 1^st^ instar nymph 3-day after hatching. (e, k, q) 2^nd^ instar nymph 0-day after molting. (f, l, r) 2^nd^ instar nymph 1-day after molting. (s) EdU-positive cell counts per symbiotic organ at the different developmental stages. (t) H3P antibody-positive cell counts per symbiotic organ at the different developmental stages. In (s, t), different alphabetical letters indicate statistically significant differences (Steel-Dwass test, *P* < 0.05; n = 4 each).

### Unique arrangement of visceral muscles in the symbiotic organ

Next, we examined the patterns of visceral muscle fibers in the nymphal midgut symbiotic organ. Previous studies reported that, in diverse insects, the midgut is surrounded by two types of muscle fibers, circular muscles running inside and longitudinal muscles running outside (**Fig. 3a**) (20). These two types of muscle fibers were certainly observed in the nymphal midgut of *P. stali* (**Fig. 3b**). When the arrangement of the visceral muscle fibers in the midgut of 2^nd^ instar nymphs was visualized using fluorochrome-labeled phalloidin, the M1, M2 and M3 regions exhibited the typical lattice-like arrangement of the muscle fibers (**Fig. 3c-e**). By contrast, the symbiotic M4 region exhibited a unique muscular pattern. While the longitudinal muscle fibers were distributed evenly, the circular muscle fibers were concentrated at the border of the crypts as if they delineate the crypt boundaries (**Fig. 3f**). TEM observations confirmed that, in the process of crypt formation, the circular muscle fibers are located at the crypt boundaries (**Fig. 3g, h**).

**FIG 3.**
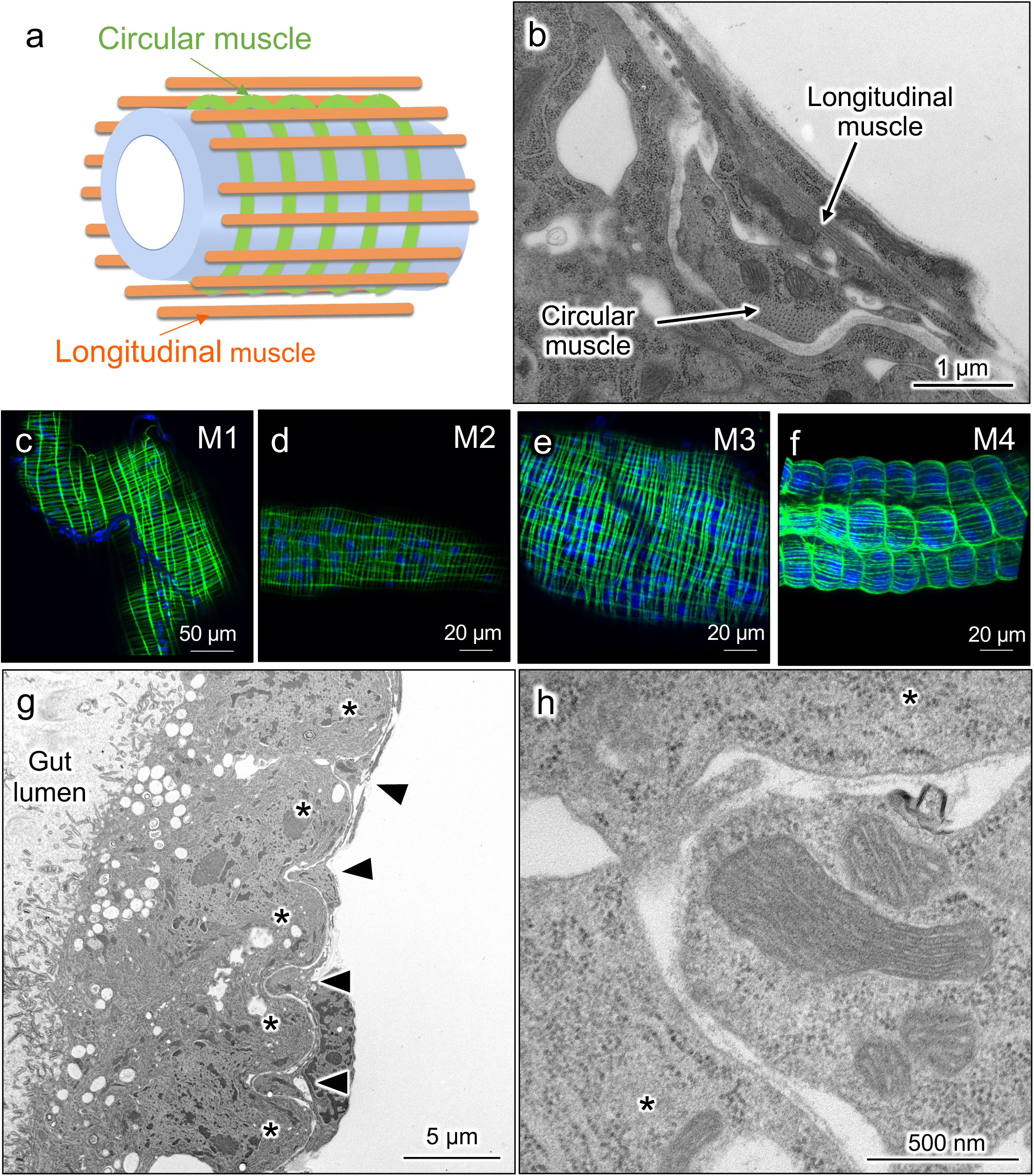
Arrangement of visceral muscle fibers on the nymphal alimentary tract of *P. stali*. (a) A schematic diagram displaying the arrangement of visceral muscles on the insect midgut. Green and orange show circular muscles and longitudinal muscles, respectively, while blue tube shows intestinal epithelium. (b) A transmission electron micrograph of the longitudinal section of the outer surface of the midgut in 1^st^ instar nymph, on which the circular muscle and the longitudinal muscle are seen. (c-f) Visualization of visceral muscle fibers on the midgut M1 region (c), M2 region (d), M3 region (e) and symbiotic M4 region (f) in late (3-day) 2^nd^ instar nymphs. Actin fibers (green) and cell nuclei (blue) are visualized by phalloidin staining and DAPI staining, respectively. Note that typical patterns of circular and longitudinal muscles are seen in M1, M2 and M3, whereas atypical patterns are observed in M4. (g) Longitudinal section of M4 region of 1^st^ instar nymph. Asterisks show crypts and arrow heads show circular muscles. (h) Enlarged image of (g). Muscle fibers are seen between crypts.

### Observation of visceral muscles in the symbiotic organ immediately after hatching

In order to understand how such peculiar muscular patterns are formed in the nymphal symbiotic organ, we observed the visceral muscles in the alimentary tract of newborn nymphs of *P. stali* immediately after hatching. Interestingly, we found that the muscular patterns of the symbiotic M4 region were already distinct from the typical lattice-like muscular patterns as observed in the M1, M2 and M3 regions (see **Fig. 3c-e**). Observation of the phalloidin-stained M4 preparations from different directions (**Fig. 4a-c**) revealed that, specifically in the symbiotic M4 region, the circular muscles exhibited characteristic bifurcated and/or curved patterns (**Fig. 4d-f**). Based on these observations, we conceived that, already in the newborn nymphs, the bifurcated circular muscles define the intestinal epithelial areas arranged in two rows, which may represent a prepattern relevant to the spatial arrangement of the crypts (**Fig. 4g**).

**FIG 4.**
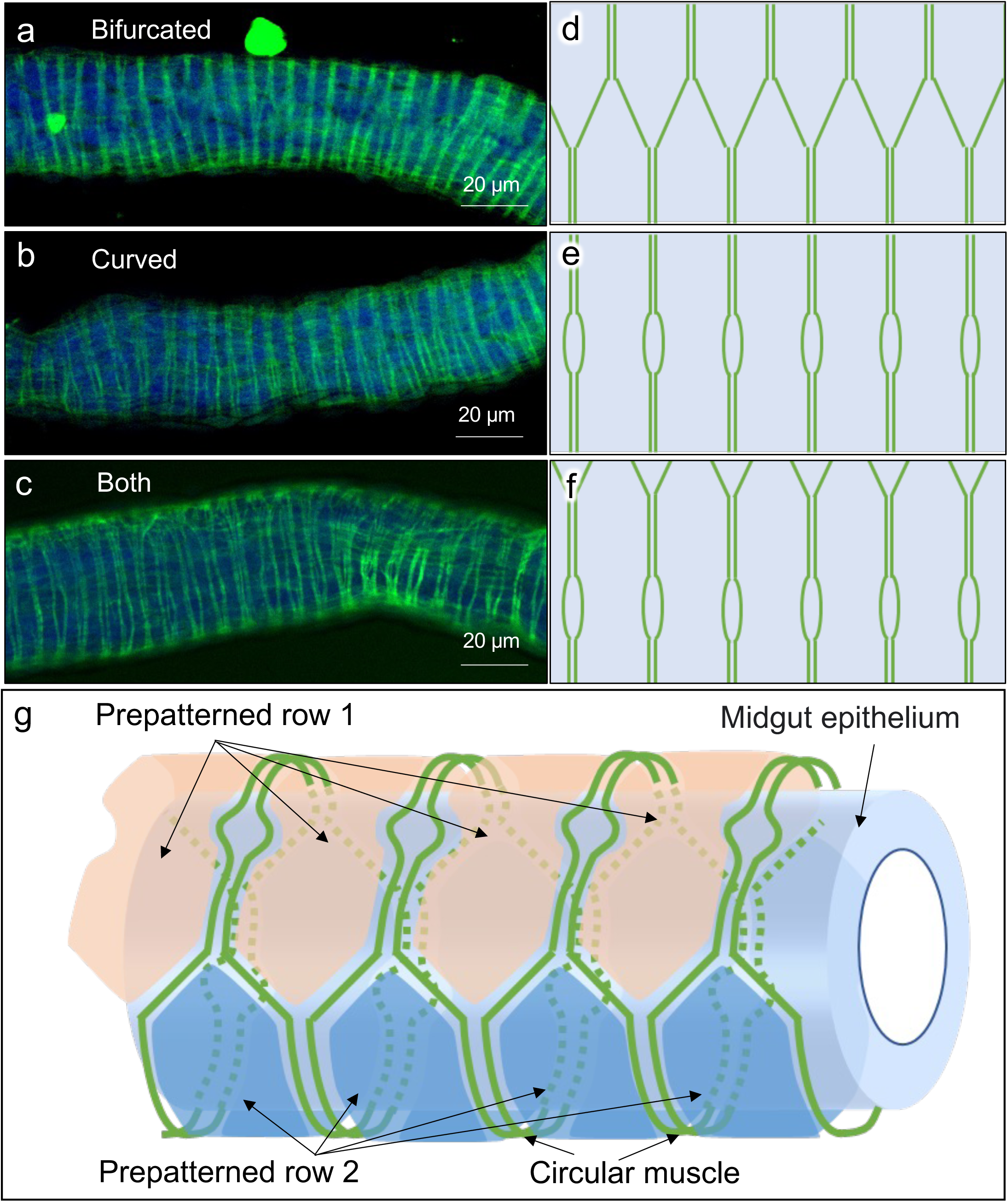
Arrangement of visceral muscle fibers on the midgut symbiotic organ of newborn nymphs of *P. stali*. (a-c) Three typical patterns of visceral muscle arrangement observed on the symbiotic organ of newborn nymphs without crypts. (a) Circular muscles are bifurcated, connected, and arranged alternately. (b) Circular muscles are curved and separated locally. (c) Both the bifurcated patterns and the curved patterns are observed simultaneously. In (a-c), actin fibers (green) and cell nuclei (blue) are visualized by phalloidin staining and DAPI staining, respectively. (d-f) Schematic diagrams of the arrangement of the circular muscle fibers corresponding to (a-c). (g) Schematic representation of the arrangement of the circular muscle fibers on the midgut symbiotic organ of newborn nymphs. The bifurcated circular muscles (green lines) delineate the intestinal epithelium into the areas arranged alternately in two prepatterned rows (orange and blue areas), although the patterns are morphologically unrecognizable at this developmental stage.

### Observation of visceral muscle arrangement during embryogenesis

How is the peculiar pattern of the visceral muscles in the newborn nymphs formed? In order to address this question, we inspected the formation process of the visceral muscles during the embryogenesis of *P. stali*. From the upper side of the eggs, embryonic eyes and egg tooth were visible 3-day after oviposition and on (**Fig. S1a**). Coincident with this, the embryonic alimentary tract was recognizable from 3-day after oviposition and on (**Fig. S1b-e**). In early 3-day embryos, neither the midgut regions nor the visceral muscle fibers were evident in the tiny alimentary tract (**Fig. S1b, f**). In late 3-day embryos, the alimentary tract became larger and developed two constrictions, which presumably corresponded to the borders of the M2, M3 and M4 regions (**Fig. S1c, g**), whereas visceral muscle fibers were not clearly recognizable (**Fig. S2a-c**). In 4-day and 5-day embryos, the midgut M1, M2, M3 and M4 regions were differentiated, in which the patterns of the visceral muscle fibers were seen (**Fig. S1d, e, h, i; Fig. S2d-f**). These observations indicated that the characteristic visceral muscle pattern in the symbiotic M4 region is formed during the embryonic development.

### Observation of visceral muscle arrangement during the process of crypt formation

As described above, immediately after hatching, the midgut epithelium of the symbiotic M4 region was divided into the areas arranged in two rows by the circular muscles (see **Fig. 4g**). On the other hand, after the crypts were formed, the circular muscles were located at the bases of the crypts arranged in four rows (see **Fig. 3f**). How, then, is the prepattern of two epithelial rows transformed into the arrangement of four crypt rows? In order to address this question, we inspected the formation processes of the visceral muscle fibers as well as the crypts in the symbiotic M4 region from 1^st^ to 3^rd^ nymphal instar of *P. stali* (**Fig. 5**). We observed the phalloidin-labeled symbiotic M4 region in detail, particularly focusing on the following three directions (also see **Fig.4a-f**): the angle from which the circular muscles are bifurcated (**Fig. 5a, d, g, j, m, p**); the angle from which the circular muscles are curved (**Fig. 5b, e, h, k, n, q**); and the angle from which both the bifurcated and curved muscles are seen (**Fig. 5c, f, i, l, o, r**). During the 1^st^ nymphal instar, the visceral muscles exhibited substantially similar patterns that delineated the epithelial areas into two rows (**Fig. 5a-i**), while primordial crypts started to form and some muscle fibers looked to accumulate between them in late 1^st^ instar (**Fig. 5g-i**). In early 2^nd^ instar nymphs, notably, crossing muscle fibers became evident to connect adjacent circular muscles at the curved sites (arrowheads in **Fig. 5k, l**). The connecting muscle fibers gradually became thicker (**Fig. 5n, o**) and finally separated each of the midgut epithelial areas of the symbiotic M4 region (**Fig. 5q, r**). FIB-SEM tomography provided high-resolution images as to how the crossing muscle fiber was formed to connect the adjacent circular muscles that delineated the crypts of the nymphal symbiotic organ (**Fig. 6; Movie S1**). In this way, the epithelium of the symbiotic M4 region was divided into four rows of crypt-to-be areas by two types of crossing muscle fibers: the bifurcated circular muscles that already existed at the time of hatching, and the connecting muscle fibers that are newly formed in parallel with the formation of the crypts.

**FIG 5.**
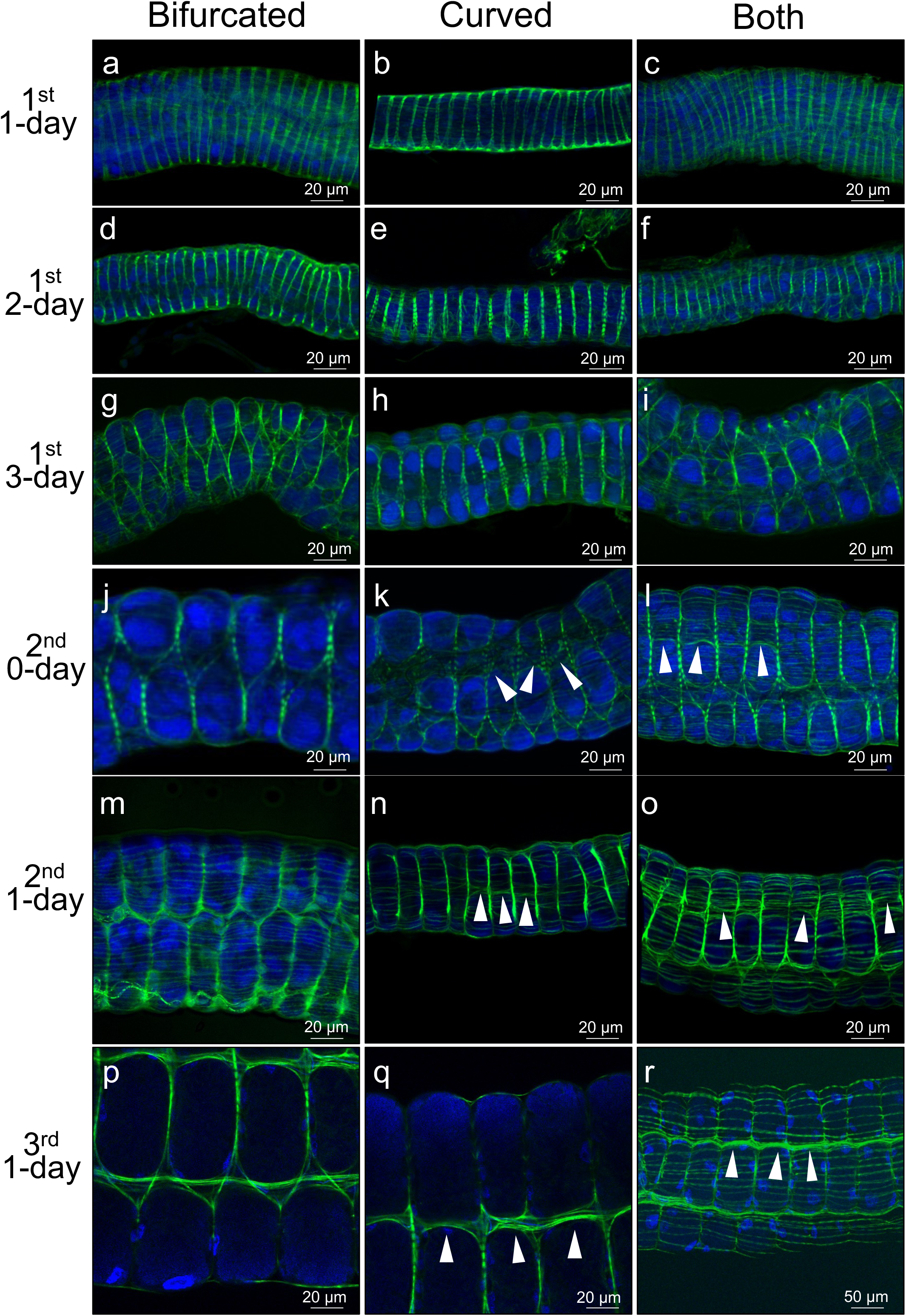
Arrangement of visceral muscle fibers during the crypt morphogenesis in the nymphal development of *P. stali*. (a-c) 1^st^ instar nymph 1-day after hatching. (d-f) 1^st^ instar nymph 2-day after hatching. (g-i) 1^st^ instar nymph 3-day after hatching. (j-l) 2^nd^ instar nymph 0-day after molting or 4-day after hatching. (m-o) 2^nd^ instar nymph 1-day after molting or 5-day after hatching. (p-r) 3^rd^ instar nymph 1-day after molting or 9-day after hatching. (a, d, g, j, m, p) Showing the patterns in which the circular muscles are bifurcated. (b, e, h, k, n, q) Showing the patterns in which the circular muscles are curved. (c, f, i, l, o, r) Showing the patterns in which both the bifurcated and curved sites are seen. At the time point of 2^nd^ instar molt and on, crossing muscle fibers that connect adjacent circular muscles are newly formed (arrowheads). Actin fibers (green) are visualized by phalloidin staining whereas cell nuclei (blue) are visualized by DAPI staining.

**FIG 6.**
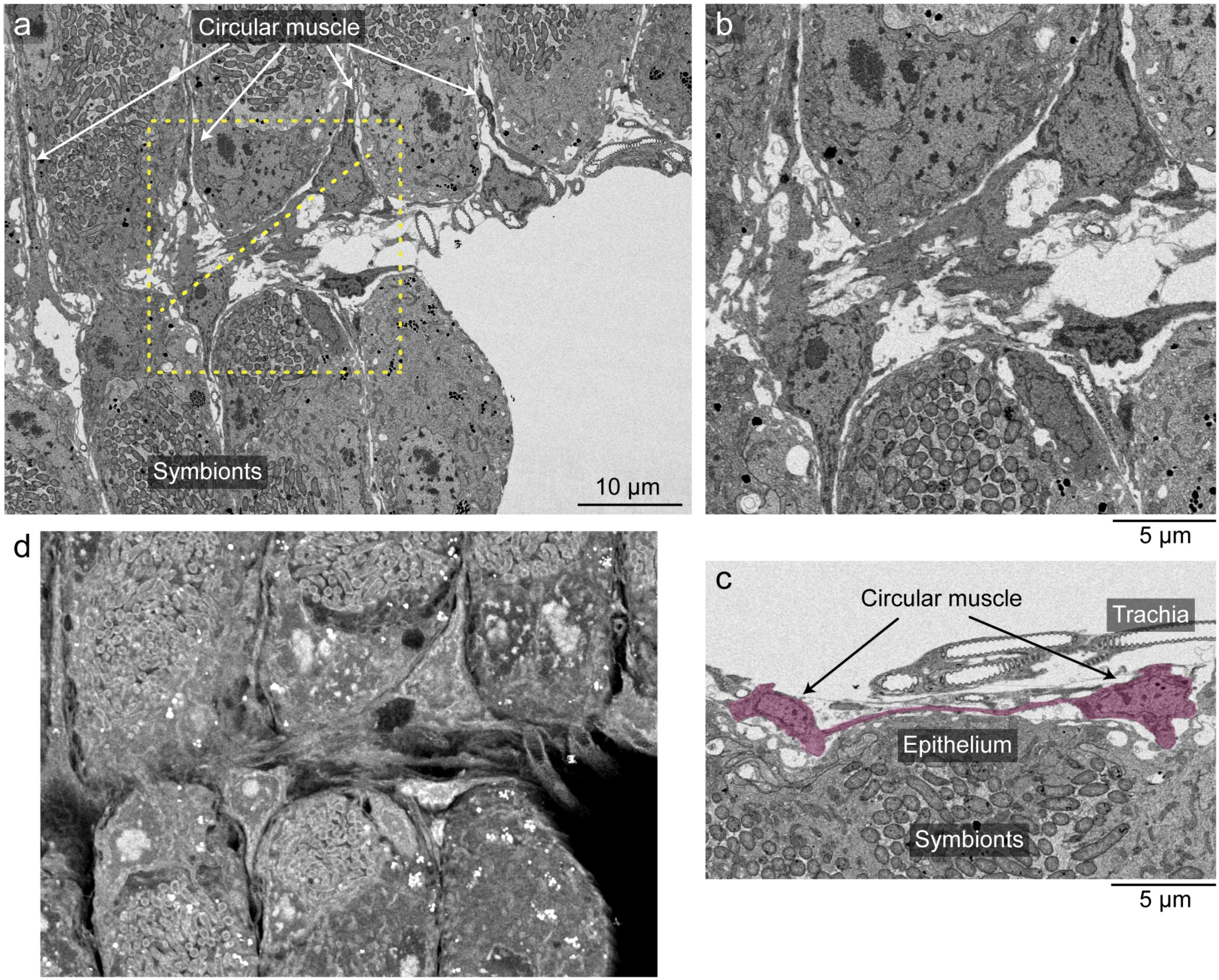
FIB-SEM tomography of the newly-formed muscle fibers connecting adjacent circular muscles in the nymphal symbiotic organ of *P. stali*. (a) Single cross-section (xy-direction) image of the midgut M4 region of 2^nd^ instar nymph 1-day after molting. Muscle fibers bridging the adjacent circular muscles is highlighted in the dotted box. (b) Magnified image of the dotted box in (a). (c) Z-directional reconstructed image of the dotted line region in (a), in which a thin cellular projection connecting two adjacent circular muscle cells is highlighted in red. (d) Z-directional reconstructed 3D image of the dotted box region in (a). See also **Movie S1**.

### Cell proliferation, crypt formation, and visceral muscle arrangement in the symbiotic organ of symbiont-free nymphs

Considering the close relationship between the symbiotic organ and the symbiotic bacteria in *P. stali*, it seemed plausible that the symbiotic bacteria may affect or induce the development of the symbiotic organ. In order to test this hypothesis, we experimentally generated symbiont-free nymphs of *P. stali* by egg surface sterilization (16, 17), and investigated the process of cell proliferation, crypt formation, and visceral muscle arrangement in the symbiotic M4 region during the early development of the aposymbiotic nymphs (**Fig. 7**). Strikingly, even in the absence of the symbiotic bacteria, similar patterns of cell proliferation (**Fig. 7a-n**), crypt formation and visceral muscle arrangement (**Fig. 7o-t**) were observed in comparison with the normal nymphs infected with the symbiotic bacteria (see **Figs. 2, 4, 6**), although growth and survival of the aposymbiotic nymphs were severely suppressed in the 2^nd^ instar and on due to the absence of the indispensable symbiont (16, 17). These results uncovered that morphogenesis of the symbiotic organ in *P. stali* proceeds autonomously even in the absence of the symbiotic bacteria.

**FIG 7.**
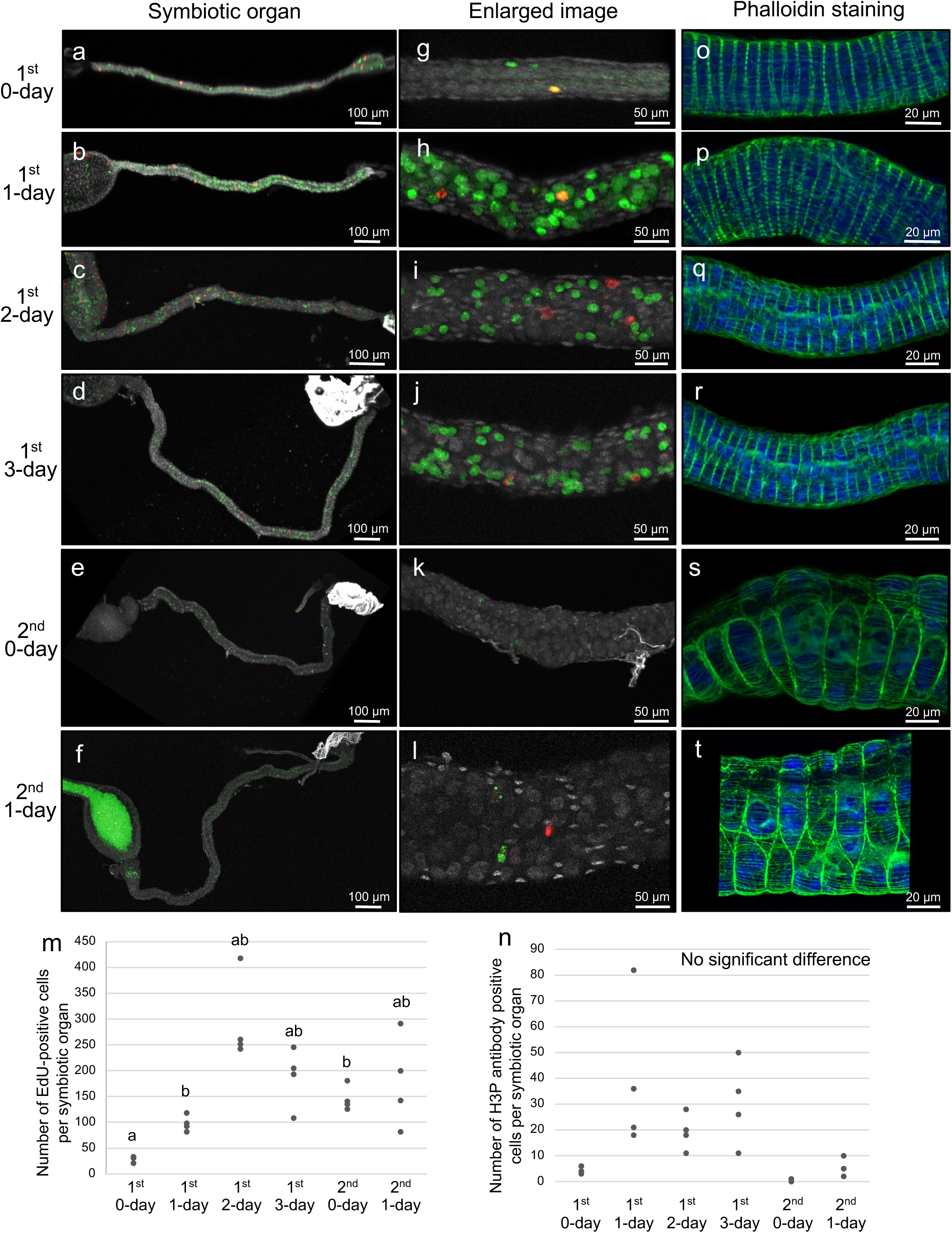
Cell proliferation, crypt formation, and visceral muscle arrangement in the symbiotic organ of symbiont-free nymphs of *P. stali*. Symbiont-free nymphs of *P. stali* were generated by egg surface sterilization as described previously (16). (a-f) Visualization of DNA synthesizing cells by EdU labeling (green), dividing cells by H3P antibody staining (red), and all intestinal cells by DNA staining (white) in dissected midgut preparations. (g-l) Enlarged images corresponding to (a-f) at the midgut symbiotic region where crypts are formed. (m) EdU-positive cell counts per symbiotic organ at the different developmental stages. (n) H3P antibody-positive cell counts per symbiotic organ at the different developmental stages. In (m, n), different alphabetical letters indicate statistically significant differences (Steel-Dwass test, *P* < 0.05, n = 4 each). (o-t) Visualization of muscle fibers by phalloidin staining (green) and cell nuclei by DAPI staining (blue) at the midgut symbiotic region where crypts are formed. (a, g, o) 1^st^ instar nymph 0-day after hatching. (b, h, p) 1^st^ instar nymph 1-day after hatching. (c, i, q) 1^st^ instar nymph 2-day after hatching. (d, j, r) 1^st^ instar nymph 3-day after hatching. (e, k, s) 2^nd^ instar nymph 0-day after molting. (f, l, t) 2^nd^ instar nymph 1-day after molting. The intense green signal in (f) is autofluorescence of food material.

## DISCUSSION

In this study, we investigated the developmental and morphogenetic processes of the midgut symbiotic organ of *P. stali* in detail. We found that, in the mid to late 1^st^ instar period when the crypt formation occurs, cell proliferation is activated and visceral muscles exhibit peculiar spatial patterns that delineate crypt boundaries in the posterior midgut region where the crypt formation occurs. On the basis of these observations, we propose a hypothetical model as to how the crypts arranged in four rows are formed in the posterior midgut region of *P. stali* (**Fig. 8**). The model assumes that the peculiar pattern of the circular muscles and the newly formed muscular bridges in the nymphal posterior midgut define the crypt boundaries (**Fig. 8a**), and the activated proliferation of the intestinal epithelial cells constrained by the muscular fiber network results in the formation of crypts as protruding epithelial outgrowths arranged in four rows (**Fig. 8b**). To confirm this hypothesis, experimental manipulation and disturbance of cell proliferation and muscle fiber formation in the nymphal posterior midgut, which can be performed either by genetic engineering or by pharmaceutical treatment, will provide important clues to understanding the mechanisms underpinning the crypt formation.

**FIG 8.**
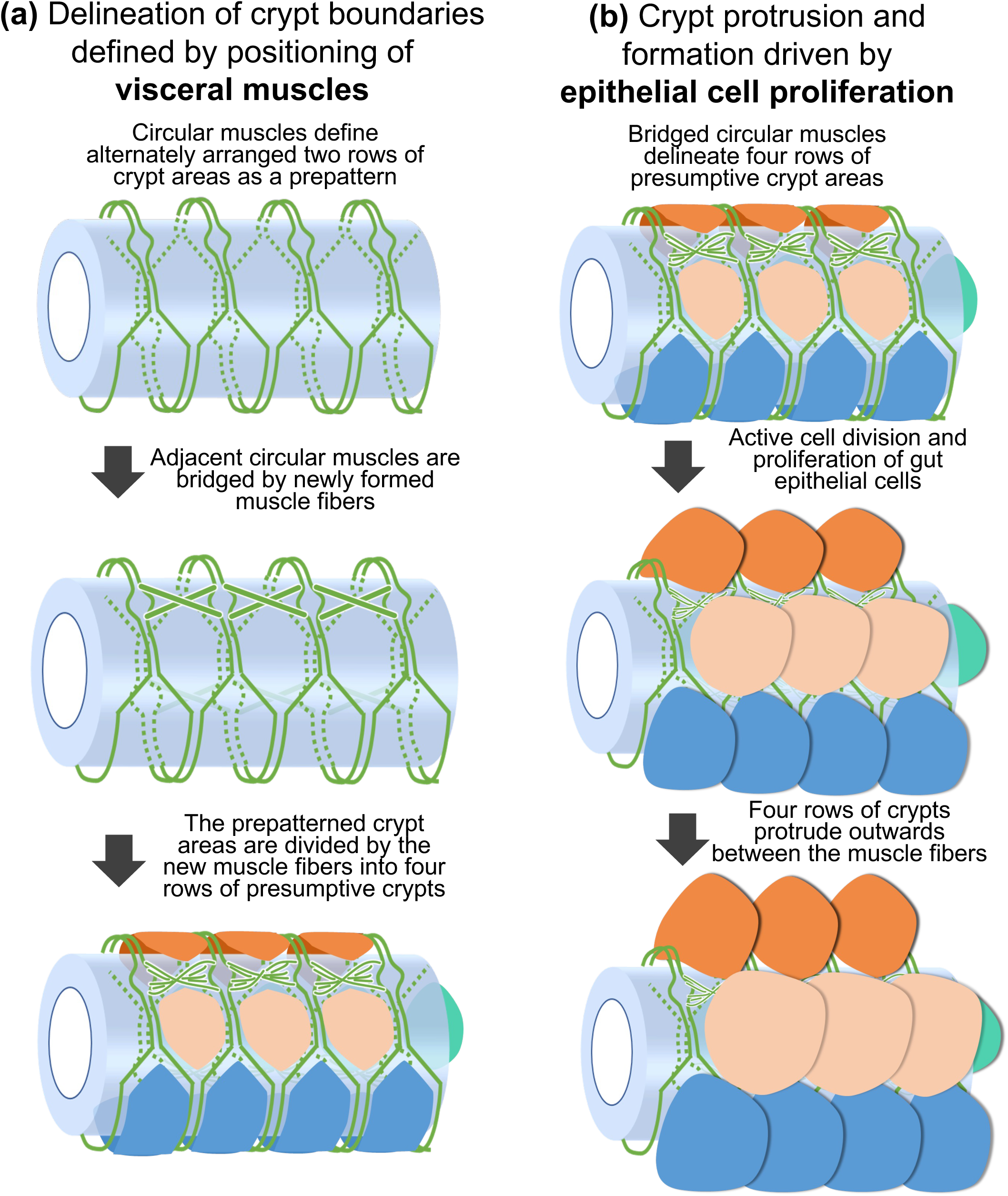
Hypothetical model of crypt morphogenesis in the symbiotic organ of *P. stali*. (a) Delineation of crypt boundaries defined by positioning of visceral muscles. (b) Crypt protrusion and formation driven by epithelial cell proliferation. The processes (a) and (b) proceed in this order with some overlap, thereby forming crypts arranged in four rows in the posterior midgut of *P. stali*.

In some symbiotic systems, microbe-derived factors are important for induction and morphogenesis of host’s symbiotic organs. For example, *Rhyzobium*-derived lipochito-oligosaccharides, called nod factors, trigger and induce the morphogenesis of leguminous plant roots into the symbiotic organs, known as root nodules, for symbiotic nitrogen fixation (21). However, we found that the crypt formation proceeds not only in normal symbiotic nymphs but also in aposymbiotic nymphs of *P. stali* (**Fig. 7**), indicating that the morphogenesis of the symbiotic organ proceeds autonomously even in the absence of the symbiotic bacteria. Hence, it is suggested that symbiont-derived factors are not needed for inducing the crypt formation. Plausibly, the crypt formation is deeply implemented into the normal development of *P. stali*, which must reflect the presumably ancient evolutionary origin of the midgut symbiotic organ conserved among diverse stinkbugs (1, 9, 22). Similarly, persistence of the symbiotic organs when deprived of their microbial symbionts has been reported in *Cassida* leaf beetles (23), *Oryzaephilus* saw-toothed grain beetles (24), and others (1).

Here it should be noted that, while diverse stinkbugs commonly develop crypts in the posterior midgut for hosting their specific symbiotic bacteria, their structural configuration exhibits considerable diversity across the stinkbug groups: four rows of crypts in Pentatomidae and Scutelleridae (19, 25); two rows of crypts in Acanthosomatidae, Coreidae, Alydidae and other families (22, 26); and ring-shaped folds like a vacuum hose in Urostylididae, Plataspidae and other families (27, 28). The formation process of the visceral muscle network in the early nymphal development of these stinkbug groups is of great interest, which should be pursued in future studies. In this context, inspection of the visceral muscle network in stinkbugs of the genus *Arma* will be of particular interest, on the ground they belong to Pentatomidae but lack the midgut crypts probably because of their carnivorous lifestyle exceptional in Pentatomidae (1).

Besides the stinkbugs, a variety of insects develop multiple blind sac-like structures, called gastric caeca or crypts, associated with the alimentary tract and use them as specialized organs for hosting their microbial symbionts, as in tortoise leaf beetles (23, 29, 30), reed beetles (31, 32), anobiid drugstore beetles (33, 34), olive flies (1, 35) and others. Originally, such gut-associated pouch-like structures must have been not for symbiosis but for increasing the inner surface area of the alimentary tract for facilitating digestion and absorption (36). In several model insects, molecular, cellular and endocrinological mechanisms underpinning the development of such gut-associated structures have been investigated. In the red flour beetle *Tribolium castaneum*, the morphogenesis of midgut-associated pouches upon metamorphosis is regulated by 20-hydroxyecdysone (37). In the yellow mealworm beetle *Tenebrio molitor*, actively dividing presumptive stem cells in the midgut epithelium form regularly distributed pouches protruding to the hemocoel after adult molting, where the regular distribution patterns are suspected to emerge via lateral inhibition (38). In the fruit fly *Drosophila melanogaster*, the development of four gastric caeca located at the foregut-midgut junction is affected by such transcription factors and morphogens as *Sex combs reduced*, *Ultrabithorax*, *Antennapedia*, *labial*, *decapentaplegic*, and *wingless*, and also matrix metalloproteinases and autophagy (39–42). Whether these mechanisms are also involved in the development and morphogenesis of the symbiotic organ in *P. stali* is of interest and to be established in future studies.

In the stinkbug *P. stali*, we found that the spatial arrangement of the visceral muscle fibers is likely involved in the morphogenesis of the symbiotic organ. Notably, though apart from insects, previous studies reported interesting cases in which visceral muscular patterns govern the morphogenesis of internal organs in vertebrates. In chick, there are numerous villi in the small intestine, and it was shown that the visceral muscles play important roles in the villus formation during embryogenesis (43). Morphogenesis of mammalian lungs entails highly regulated terminal branching of tracheae, and it was reported that muscle fibers define the patterning of the tracheal bifurcation during murine embryogenesis (44, 45). Needless to say, the morphogenesis and patterning mediated by muscle fiber delineation must have evolved independently in insects and vertebrates, but we point out that similar developmental and morphogenetic mechanisms may be operating more widely than previously envisioned.

In conclusion, we uncovered previously unrecognized muscular patterning and cell proliferation in the early nymphal midgut epithelium that are presumably responsible for the characteristic morphogenesis of the symbiotic organ of the stinkbug *P. stali*, which shed light on how such intimate host-microbe mutualistic associations are initiated and established. What molecular and cellular mechanisms underpin the muscular patterning and cell proliferation are of great interest, which we are challenging to elucidate by RNA sequencing of the dissected early posterior midgut samples, picking up specifically up-regulated genes at the morphogenetic stage, and RNAi knockdown of the candidate genes to identify the genes involved in the morphogenesis of the midgut symbiotic organ.

## MATERIALS AND METHODS

### Insect material

In this study, we used a mass-reared laboratory strain of *P. stali*, which had been established from adult insects collected in Tsukuba, Ibaraki, Japan. Insect rearing was conducted essentially as described previously (16, 19, 46). In each plastic container (15 cm in diameter, 5 cm high), 10-20 adult insects were fed with raw peanuts and distilled water (DW) containing 0.05% ascorbic acid (DWA), and allowed to lay eggs on filter paper placed inside the container. An egg mass of *P. stali* usually consists of 14 eggs. Each egg mass was transferred to a plastic Petri dish (8.5 cm in diameter, 2 cm high) in which filter paper on the bottom, three grains of raw peanuts and DWA were supplied (47). We inspected egg hatching and nymphal molting in the rearing containers every day between 12:00 and 17:00, and renewed peanuts and DWA once a week.

### Cytological visualization of muscle fibers

The insects were dissected under a stereomicroscope (SZ61, Olympus, Japan) in phosphate buffered saline (PBS: 137 mM NaCl, 8.1 mM Na_2_HPO_4_, 2.7 mM KCl, pH = 7.4) using tweezers, razor and scissors. The isolated tissues were fixed in buffered PFA (PBS containing 4% paraformaldehyde) and thoroughly washed with PBST (PBS containing 0.1% Tween 20). After incubated in PBS, the tissue samples were stained with Alexa Fluor™ 488 Phalloidin (Invitrogen, USA), washed thoroughly with PBS, subjected to DNA staining with PBS supplemented with 1 µg/mL 4’,6-diamidino-2-phenylindole dihydrochloride (DAPI), and observed under a fluorescence stereomicroscope (M165FC, Leica, Germany) and a confocal laser scanning microscope (LSM700, Zeiss, Germany).

### Cytological visualization of proliferating and dividing cells

We performed cell proliferation analysis of the symbiotic organ using 5-ethynyl-2’-deoxyuridine (EdU), a thymidine analog that is incorporated into newly synthesized DNA, and phosphorylated Histone H3 (H3P) antibody that binds to dividing cells specifically. The insects were injected with 50 nL of 10 mM EdU solution, and 1 h after injection, the midgut symbiotic organs were dissected and fixed with buffered PFA for 1 h. The fixed tissue samples were subjected to fluorescence labeling of EdU using Click-iT EdU Imaging Kits Alexa Flour 488 (Invitrogen, USA). Subsequently, the tissue samples were treated with blocking buffer (PBS supplemented with 1% bovine serum albumin), incubated with anti-Histone H3S10ph antibody (rabbit, polyclonal, GeneTex, USA) for 30 min, and incubated with anti-rabbit fluorescent antibody (goat, monoclonal, Alexa Fluor 555) for 30 min. Then, the tissue samples were counter-stained with PBS supplemented with 1 µg/mL DAPI, and observed under a confocal microscope (LSM700, Zeiss, Germany).

### Fluorescence in situ hybridization (FISH)

FISH was conducted using a fluorochrome-labeled oligonucleotide probe SymAC89R (5’-Alexa Fluor 555-GCA AGC TCT TCT GTG CTG CC-3’) that targeted bacterial 16S rRNA of the symbiont as described previously (19, 48). Dissected symbiotic organs were fixed in buffered PFA for 3 h at room temperature, and washed with PBST. For whole-mount FISH, the organs were washed and equilibrated with a hybridization buffer (20 mM Tris-HCl [pH 8.0], 0.9 M NaCl, 0.01% sodium dodecyl sulfate, 30% formamide) and then incubated in the hybridization buffer supplemented with 100 nM probe and 1 μg/ml DAPI overnight at room temperature in a dark box. After the incubation, the samples were washed with PBST, mounted with 80% glycerol or Slowfade Gold Antifade Mountant (Thermo Fisher, USA), and observed under a fluorescence stereomicroscope (M165FC, Leica, Germany) or a confocal microscope (LSM700, Zeiss, Germany).

### Transmission electron microscopy (TEM)

Nymphs of *P. stali* were staged and their abdominal parts containing the almost entire midgut were dissected in cold PBS to remove their ventral cuticles. The samples were prefixed overnight at 4°C in the mixture of 4% PFA (Electron Microscopy Sciences, 15710) and 2% glutaraldehyde (Sigma-Aldrich, G5882) in PBS, postfixed with 2% osmium tetroxide in DW at 4°C for 2 h, dehydrated in a graded ethanol series (50%, 60%, 70%, 80%, 90%, 95%, 99%, and 100%), and treated with 100% propylene oxide followed by Epon 812. From each of the samples, the M4 region was isolated by forceps and embedded in Epon 812. Ultrathin sections of 70-nm thickness were cut with a diamond knife, collected on single-hole grids with a support membrane, stained with uranyl acetate and lead citrate, and observed with a transmission electron microscope (Hitachi, H-7650).

### Focused ion beam scanning electron microscope (FIB-SEM) tomography

The samples for FIB-SEM were prepared by a modified Ellisman method (49). Nymphs of *P. stali* were dissected and prefixed as described above, and incubated with 1.5% potassium ferrocyanide followed by 2% osmium tetroxide in DW at 4°C for 2 h. The tissues were washed with DW and postfixed with 2% osmium tetroxide in DW at room temperature (RT) for 30 min. After washing with DW, the specimens were then stained en bloc in a solution of 4% uranyl acetate dissolved in DW overnight for contrast enhancement and then washed with DW. Subsequently, the specimens were further stained with Walton’s lead aspartate solution for 30 min at 60°C. After washing with DW, the specimens were dehydrated and embedded in Epon 812 as described above. The trimmed specimens were set on the stage of FIB-SEM (Zeiss, Crossbeam 540). By using the Smart FIB software (Zeiss), the surface of the specimen was milled with a gallium ion FIB at 30 kV with a current of 7 nA at the pitch of 30 nm/slice, and SEM images were obtained at a landing energy of 1.5 keV. The other parameters were as follows: beam current = 1 nA; image pixel size = 30 nm; resolution = 3,072 × 2,304 (92.16 μm × 69.12 μm); scan speed = 1.6 min/image. The SEM images were obtained by 1250 times of the slice-and-view processes, so that the total sliced thickness was 37.5 μm. The resultant image stacks were processed and edited by Dragonfly 3.1 software (ORS Inc.). The movie was edited by Clipchamp video editor (Microsoft Corporation). A cropped region (67.10 μm × 49.96 μm) of arbitrary thickness is shown in Fig. 6 and Movie S1.

### Preparation of symbiont-free insects

In *P. stali*, vertical symbiont transmission occurs via maternal smearing of symbiont-containing secretion onto the egg surface upon oviposition and nymphal probing of the symbiotic bacteria on the egg surface (16, 50). Therefore, symbiont-free nymphs can be easily generated by egg surface sterilization. The eggs were soaked in 4% formaldehyde for 10 min, washed with sterilized water for 30 min, and air-dried. Newborn nymphs from these sterilized eggs were free of the symbiotic bacteria. The symbiont-free nymphs were reared as described previously (17).

## SUPPLEMENTAL MATERIALS

FIG S1 FIG S2

Movie S1

## ACKNOWLEDGMENTS

We thank Tomoko Matsushita and Yumi Kanazawa for insect maintenance and supply, and Haruyasu Kohda and Tatsuya Katsuno for their support in electron microscopic analyses.

This study was supported by the Japan Science and Technology Agency (JST) ERATO grant no. JPMJER1902 to T.F. and T.H., the Japan Society for the Promotion of Science (JSPS) KAKENHI grant nos. JP17H06388 to T.F. and JP22K19352 to T.H., the Hakubi Project grant of Kyoto University to T.H., and Nagase Science and Technology Foundation grant to T.H. S.O. was supported by the JSPS Research Fellowships for Young Scientists no. JP20J21460.

## FIGURE CAPTIONS

**FIG S1** Embryonic development of the alimentary tract of *P. stali*. (a) Eggs after 0-day, 1-day, 2-day, 3-day, 4-day and 5-day after oviposition. The eggs of *P. stali* usually hatch in five days after oviposition under our rearing condition (19). From the upper side of the eggs, eyes and an egg tooth become visible from 3-day after oviposition and on, during which the development of the alimentary tract proceeds. (b-e) Embryonic alimentary tracts dissected from the developing eggs. (f-i) Enlarged images of the posterior end region of the alimentary tract. (b, f) In early 3-day embryos, the hindgut and the Malphigian tubules are formed, whereas the midgut is still rudimentary. Muscle fibers are still unrecognizable. (c, g) In late 3-day embryos, the midgut exhibits constrictions, presumably reflecting differentiation of midgut regions. Some muscle fibers start to form. (d, h) In 4-day embryos, the midgut M1, M2, M3 and M4 regions are formed, with muscle fibers evidently seen. In the M4 region, circular muscles become evident. (e, i) In 5-day embryos prior to hatching, morphogenesis of the alimentary tract almost completes, with circular and longitudinal muscle fibers well-developed in the M1, M2 and M3 regions. In the M4 region, the bifurcating patterns of circular muscles, as observed in newborn nymphs (see **Fig. 3**), are seen. In (b-i), actin fibers (green) and cell nuclei (blue) are visualized by phalloidin staining and DAPI staining, respectively.

**FIG S2** Development of visceral muscle fibers during embryogenesis of *P. stali*. (a-c) Actin fibers (green) and nuclear DNA (blue) visualized in M2 (a), M3 (b) and symbiotic M4 (c) regions of 3-day embryos. Visceral muscles are obscure. (d-f) Actin fibers (green) and nuclear DNA (blue) visualized in M2 (d), M3 (e) and symbiotic M4 (f) regions of 4-day embryos. Circular and longitudinal muscle fibers are clearly seen.

**Movie S1** FIB-SEM tomography of the newly-formed muscle fibers connecting adjacent circular muscles in the nymphal symbiotic organ of *P. stali*. Z-directional view (sequential sections from surface to interior) of the midgut M4 region of 2^nd^ instar nymph 1-day after molting. See also **Fig. 6**.

## REFERENCES

1. Buchner P. 1965. Endosymbiosis of animals with plant microorganisms. Interscience, New York.

2. Bourtzis K, Miller TA. 2003. Insect Symbiosis. CRC Press.

3. Douglas AE. 2009. The microbial dimension in insect nutritional ecology. Funct Ecol 23:38–47.

4. Brune A. 2014. Symbiotic digestion of lignocellulose in termite guts. Nat Rev Microbiol 12:168–180.

5. Flórez LV, Biedermann PH, Engl T, Kaltenpoth M. 2015. Defensive symbioses of animals with prokaryotic and eukaryotic microorganisms. Nat Prod Rep 32:904–936.

6. Lemoine MM, Engl T, Kaltenpoth M. 2020. Microbial symbionts expanding or constraining abiotic niche space in insects. Curr Opin Insect Sci 39:14–20.

7. Douglas AE. 2020. Housing microbial symbionts: evolutionary origins and diversification of symbiotic organs in animals. Philos Trans R Soc Lond B Biol Sci 375:20190603.

8. Koga R, Meng XY, Tsuchida T, Fukatsu T. 2012. Cellular mechanism for selective vertical transmission of an obligate insect symbiont at the bacteriocyte–embryo interface. Proc Natl Acad Sci USA 109:E1230–E1237.

9. Salem H, Florez L, Gerardo N, Kaltenpoth M. 2015. An out-of-body experience: the extracellular dimension for the transmission of mutualistic bacteria in insects. Proc Royal Soc B 282:20142957.

10. Gilbert SF, Bosch TC, Ledón-Rettig C. 2015. Eco-Evo-Devo: developmental symbiosis and developmental plasticity as evolutionary agents. Nat Rev Genet 16:611–622.

11. Fronk DC, Sachs JL. 2022. Symbiotic organs: the nexus of host–microbe evolution. Trends Ecol Evol 37:599–610.

12. Alarcón ME, Polo PG, Akyüz SN, Rafiqi AM. 2022. Evolution and ontogeny of bacteriocytes in insects. Front Physiol 13:1034066.

13. Braendle C, Miura T, Bickel R, Shingleton AW, Kambhampati S, Stern DL. 2003. Developmental origin and evolution of bacteriocytes in the aphid–*Buchnera* symbiosis. PLoS Biol 1:e21.

14. Matsuura Y, Kikuchi Y, Miura T, Fukatsu T. 2015. *Ultrabithorax* is essential for bacteriocyte development. Proc Natl Acad Sci USA 112:9376–9381.

15. Rafiqi AM, Rajakumar A, Abouheif E. 2020. Origin and elaboration of a major evolutionary transition in individuality. Nature 585:239–244.

16. Hosokawa T, Ishii Y, Nikoh N, Fujie M, Satoh N, Fukatsu T. 2016. Obligate bacterial mutualists evolving from environmental bacteria in natural insect populations. Nat Microbiol 1:15011.

17. Nishide Y, Onodera NT, Tanahashi M, Moriyama M, Fukatsu T, Koga R. 2017. Aseptic rearing procedure for the stinkbug *Plautia stali* (Hemiptera: Pentatomidae) by sterilizing food-derived bacterial contaminants. Appl Entomol Zool 52:407–415.

18. Moriyama M, Hayashi T, Fukatsu T. 2022. A mucin protein predominantly expressed in the female-specific symbiotic organ of the stinkbug *Plautia stali*. Sci Rep 12:7782.

19. Oishi S, Moriyama M, Koga R, Fukatsu T. 2019. Morphogenesis and development of midgut symbiotic organ of the stinkbug *Plautia stali* (Hemiptera: Pentatomidae). Zool Lett 5:16.

20. Campos-Ortega JA, Hartenstein V. 1985. The embryonic development of Drosophila melanogaster. Springer-Verlag, Berlin, Germany.

21. Geurts R, Bisseling T. 2002. *Rhizobium* Nod factor perception and signalling. Plant Cell 14:S239–S249.

22. Kikuchi Y, Hosokawa T, Fukatsu T. 2011. An ancient but promiscuous host–symbiont association between *Burkholderia* gut symbionts and their heteropteran hosts. ISME J 5:446–460.

23. Fukumori K, Oguchi K, Ikeda H, Shinohara T, Tanahashi M, Moriyama M, Koga R, Fukatsu T. 2022. Evolutionary dynamics of host organs for microbial symbiosis in tortoise leaf beetles (Coleoptera: Chrysomelidae: Cassidinae). mBio 13:e03691–21.

24. Koch A. 1936. Symbiosestudien. II. Experimentelle untersuchungen an *Oryzaephilus surinamensis* L. (Cucujidae, Coleopt.). Z Morphol Ökol Tiere 32:137–180.

25. Hosokawa T, Imanishi M, Koga R, Fukatsu T. 2019. Diversity and evolution of bacterial symbionts in the gut symbiotic organ of jewel stinkbugs (Hemiptera: Scutelleridae). Appl Entomol Zool 54:359–367.

26. Kikuchi Y, Hosokawa T, Nikoh N, Meng XY, Kamagata Y, Fukatsu T. 2009. Host-symbiont co-speciation and reductive genome evolution in gut symbiotic bacteria of acanthosomatid stinkbugs. BMC Biol 7:2.

27. Kaiwa N, Hosokawa T, Nikoh N, Tanahashi M, Moriyama M, Meng XY, Maeda T, Yamaguchi K, Shigenobu S, Ito M, Fukatsu T. 2014. Symbiont-supplemented maternal investment underpinning host’s ecological sdaptation. Curr Biol 24:2465–2470.

28. Koga R, Tanahashi M, Nikoh N, Hosokawa T, Meng XY, Moriyama M, Fukatsu T. 2021. Host’s guardian protein counters degenerative symbiont evolution. Proc Natl Acad Sci USA 118:e2103957118.

29. Stammer HJ. 1936. Studien an symbiosen zwischen käfern und mikroorganismen. 2: Die Symbiose des Bromius obscurus L. und der Cassida - Arten (Coleopt. Chrysomel.). Z Morphol Ökol Tiere 31:682–697.

30. Salem H, Bauer E, Kirsch R, Berasategui A, Cripps M, Weiss B, Koga R, Fukumori K, Vogel H, Fukatsu T, Kaltenpoth M. 2017. Drastic genome reduction in an herbivore’s pectinolytic symbiont. Cell 171:1520–1531.e13.

31. Stammer HJ. 1935. Studien an symbiosen zwischen käfern und mikroorganismen.1. Die symbiose der Donaciinen (Coleopt. Chrysomel.). Z Morphol Ökol Tiere 29:585–608.

32. Reis F, Kirsch R, Pauchet Y, Bauer E, Bilz LC, Fukumori K, Fukatsu T, Kölsch G, Kaltenpoth M. 2020. Bacterial symbionts support larval sap feeding and adult folivory in (semi-) aquatic reed beetles. Nat Commun 11, 2964.

33. Koch A. 1933. Über das Verhalten symbiontenfreier Sitodrepalarven. Biol Zbl 53:199–203.

34. Pant N, Fraenkel G. 1954. Studies of the symbiotic yeasts of the two insect species, Lasioderma serricorne F. and Stegobium paniceum L. Biol Bull 107:420–430.

35. Estes AM, Hearn DJ, Bronstein JL, Pierson EA. 2009. The olive fly endosymbiont, “*Candidatus* Erwinia dacicola,” switches from an intracellular existence to an extracellular existence during host insect development. Appl Environ Microbiol 75:7097–7106.

36. Dow JAT. 1987. Insect midgut function. Adv Insect Physiol 19:187–328

37. Parthasarathy R, Palli SR. 2008. Proliferation and differentiation of intestinal stem cells during metamorphosis of the red flour beetle, *Tribolium castaneum*. Dev Dyn 237:893–908.

38. Nardi JB, Bee CM, Miller LA. 2010. Stem cells of the beetle midgut epithelium. J Insect Physiol 56:296–303.

39. Panganiban GE, Reuter R, Scott MP, Hoffmann FM. 1990. A *Drosophila* growth factor homolog, *decapentaplegic*, regulates homeotic gene expression within and across germ layers during midgut morphogenesis. Development 110:1041–1050.

40. Staehling-Hampton K, Hoffmann FM, Baylies MK, Rushtont E, Bate M. 1994. *Dpp* induces mesodermal gene expression in *Drosophila*. Nature 372:783–786.

41. Page-McCaw A, Serano J, Santé JM, Rubin GM. 2003. *Drosophila* matrix metalloproteinases are required for tissue remodeling, but not embryonic development. Dev Cell 4:95–106.

42. Denton D, Shravage B, Simin R, Mills K, Berry DL, Baehrecke EH, Kumar S. 2009. Autophagy, not apoptosis, is essential for midgut cell death in *Drosophila*. Curr Biol 19:1741–1746.

43. Shyer AE, Tallinen T, Nerurkar NL, Wei Z, Gil ES, Kaplan DL, Tabin CJ, Mahadevan L. 2013. Villification: How the gut gets its villi. Science 342:212–218.

44. Metzger RJ, Klein OD, Martin GR, Krasnow MA. 2008. The branching programme of mouse lung development. Nature 453:745–750.

45. Kim HY, Pang M-F, Varner VD, Kojima L, Miller E, Radisky DC, Nelson CM. 2015. Localized smooth muscle differentiation is essential for epithelial bifurcation during branching morphogenesis of the mammalian lung. Dev Cell 34:719–726.

46. Koga R, Moriyama M, Onodera-Tanifuji N, Ishii Y, Takai H, Mizutani M, Oguchi K, Okura R, Suzuki S, Gotoh Y, Hayashi T, Seki M, Suzuki Y, Nishide Y, Hosokawa T, Wakamoto Y, Furusawa C, Fukatsu T. 2022. Single mutation makes *Escherichia coli* an insect mutualist. Nat Microbiol 7:1141–1150.

47. Kotaki T, Hata K, Gunji M, Yagi S. 1983. Rearing of the brown-winged green bug, Plautia stali Scott (Hemiptera: Pentatomidae) on several diets. J Appl Entomol Zool 27:63– 68.

48. Koga R, Tsuchida T, Fukatsu T. 2009. Quenching autofluorescence of insect tissues for *in situ* detection of endosymbionts. Appl Entomol Zool 44:281–291.

49. West JB, Fu Z, Deerinck TJ, Mackey MR, Obayashi JT, Ellisman MH. 2010. Structure– function studies of blood and air capillaries in chicken lung using 3D electron microscopy. Respir Physiol Neurobiol 170:202–209.

50. Abe Y, Mishiro K, Takanashi M. 1995. Symbiont of brown-winged green bug, *Plautia stali* SCOTT. Appl Entomol Zool 39:109–115.

